# Manipulation of the microRNA172 - *AP2L2* interaction provides precise control of wheat and triticale plant height

**DOI:** 10.1101/2024.08.05.606718

**Authors:** Chaozhong Zhang, Joshua Hegarty, Mariana Padilla, David M. Tricoli, Jorge Dubcovsky, Juan M. Debernardi

## Abstract

The *REDUCED HEIGHT* (*RHT*) dwarfing alleles *Rht-B1b* and *Rht-D1b* were essential in the “Green Revolution” to optimize wheat plant height and increase grain yield. However, those alleles reduce coleoptile length limiting sowing depth, which triggered the search for alternative dwarfing genes. In this study, we engineered the interaction between miR172 and *AP2L2* genes to fine-tune wheat and triticale plant height without affecting coleoptile and first-leaf length.

## Main

*RHT1* encodes a DELLA protein, which participates in the gibberellin (GA) growth-stimulating pathway**^1^**. GA-insensitive truncations of this protein are responsible for the semi-dwarf *Rht1b* alleles**^2^**. The growth-repressing effect of *Rht1b* alleles optimized plant height, reduced lodging and improved harvest index. However, those alleles also reduced above-ground biomass and coleoptile length. The later limits sowing depth and access to deeper soil moisture**^3^**. This has triggered the search for GA-sensitive dwarfing genes with fewer negative pleiotropic effects^4^.

Plant height in grasses is determined by the elongation of the peduncle and stem internodes, which is regulated by a complex genetic network. The microRNA172 (miR172) is a conserved regulator within this network**^5-6^**. In wheat, miR172 expression is induced during the reproductive transition, and regulates flowering time, plant height, and both spike and floret development by repressing the expression of *APETALA2*-like (*AP2L*) genes**^7^**. Reduction of miR172 activity in the semi-dwarf tetraploid wheat variety ‘Kronos’ (*Rht-B1b*) using a transgenic target mimicry (MIM172) approach delayed reproductive transition a few days and generated shorter plants with more compact spikes**^7^**.

Wheat has four *AP2L* genes targeted by miR172, and among them *AP2L2* and *AP2L5* regulate flowering transition, stem elongation and spike development**^8^**. Point mutations in the miR172 target site of the *AP2L* genes reduce miR172 activity and generate resistant alleles designated hereafter as *rAp2l*. An *rAp2l*-*A5* allele originated the domestication gene *Q* and the free-threshing wheats**^7^**. Additional mutations in the miR172 target site of *Q***^9^**, or in the homeolog *AP2L-D5***^10^**, result in plants with reduced height but, unfortunately, with associated spike defects**^9^**. In this study, we explore the effects of chemically induced alleles *rAp2l-A2* from tetraploid and *rAp2l-B2* from hexaploid wheat**^8^** (**Fig. S1a**) as well as multiple new CRISPR induced alleles.

The *rAp2l-A2* EMS-mutation in the semi-dwarf Kronos reduced stem length by 21%, whereas the introgression of the *rAp2l-B2* allele into Kronos or Kronos-*rAp2l-A2* backgrounds, reduced stem length by 43-45% (**Fig. S1a-c, Data S1**). We next used CRISPR-Cas9 with a gRNA specifically targeting the miR172 target site of *rAp2l-B2*, because *AP2L-A2* has a polymorphism that disrupts the gRNA target (**Fig. 1a, Fig. S2a**). We generated multiple independent CRISPR T_0_ events into the semi-dwarf Kronos (*Rht-B1b*) and a near-isogenic tall line (*Rht-B1a*), and classified them as wildtype, heterozygous or biallelic/homozygous (**Fig. S2, Data S2**). Most of the CRISPR mutations were small frameshift indels in the miR172 target site (**Fig. 1a, Fig. S2a**), located downstream of the conserved AP2 domains and close to the stop codon (**Fig. 1a**). Both in-frame and frameshift indels resulted in semi-dominant dwarfing effects, suggesting that disruptions of the reading-frame at the end of the gene have limited effects on AP2L2 activity. The dominance effect of the dwarfing *rAP2L-B2* alleles was similar in the tall-*Rht-B1a* plants (T_0_=58.4%, T_1_=57.0%, **Fig. S2b,c**) and the semi-dwarf *Rht-B1b* backgrounds (T_0_=52.2%, **Fig. S2d**).

**Figure 1.**
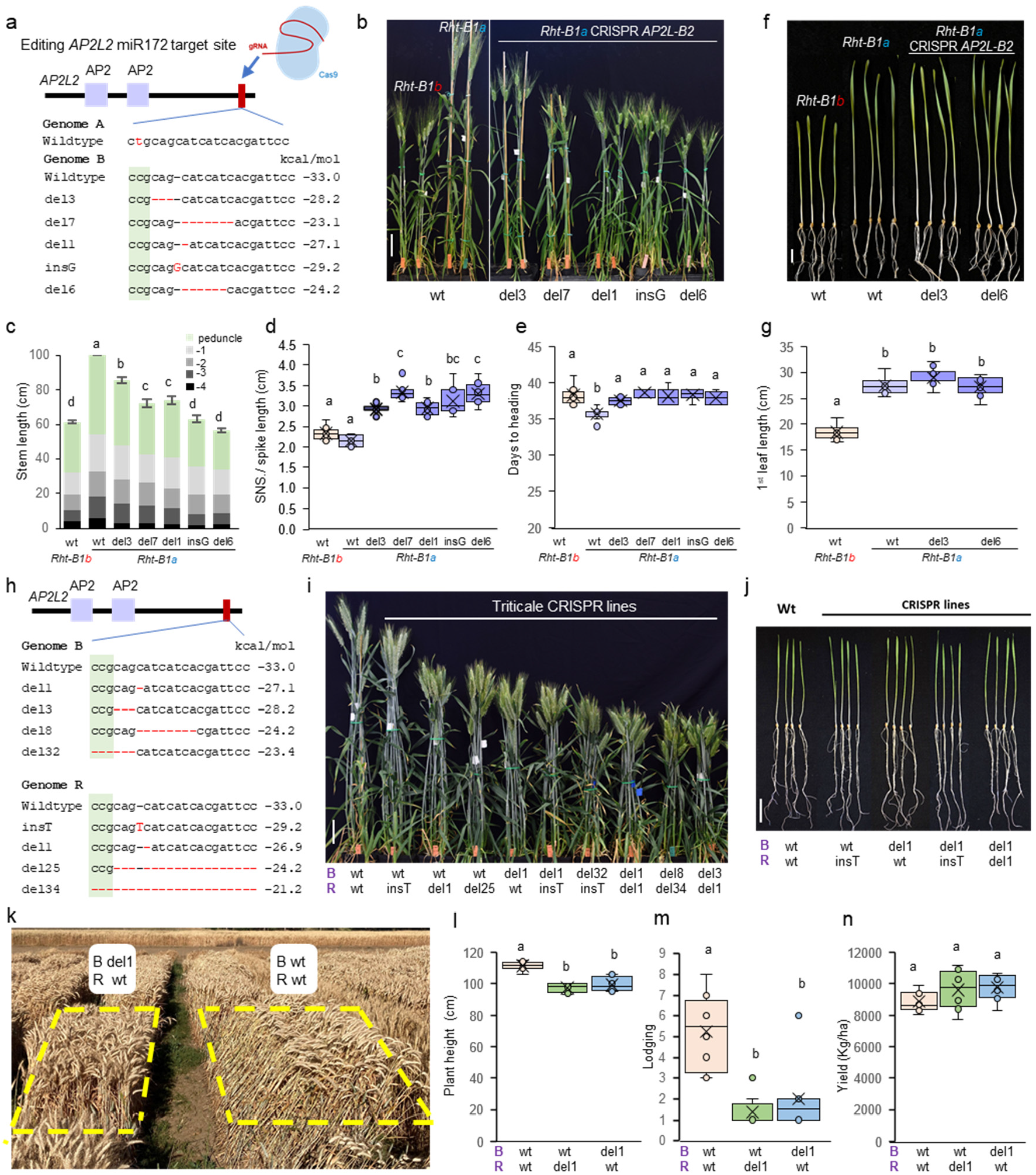
miR172-resistant *rAp2l2* alleles reduce plant height without affecting coleoptile length or yield. **a**. Schematic representation of the *AP2L2* gene indicating the AP2 domains (purple), the miR172 target site (red), and the *rAp2l-B2* alleles generated by CRISPR-Cas9 (del= deletion, ins= insertion). **b**. Kronos *Rht-B1a* and *Rht-B1b* plants, and T_2_ *rAp2l-B2* CRISPR plants 3 weeks after heading, bar = 10 cm. **c**. Stem length: internodes are in different grey colors and the peduncle is in green (n=8). **d**. Spikelet density calculated as spikelet number per spike / spike length (n=8). **e**. Days to heading (n=8). **f**. Seedlings 14 days after germination, bar = 2 cm. **g**. Length of the first leaf in 14 days-old seedlings (n=9-11). **h**. *rAp2l-B2* and *rAp2l-R2* alleles generated by CRISPR-Cas9 in triticale variety UC-Bopak. **i**. Triticale wildtype and *rAp2l2* CRISPR plants 3 weeks after heading, bar = 10 cm. The genotypes of *AP2L-B2* (B) and *AP2L-R2* (R) homeologs are indicate below each plant. **j**. Seedlings 10 days after germination, bar = 1 cm. **k-n**. Field experiment comparing triticale wildtype and CRISPR lines (n=8). **l**. Plant height. (**m**) Lodging (1-9 scale, 1= no lodging and 9=100% lodging). (**n**) Grain yield (kg/ha). Different letters above bars and plots indicate significant differences based on Tukey tests (*P*<0.05). Estimated interactions energies are indicated to the right of the sequences (in kcal/mol). Raw data and statistics are presented in **Data S3**.

We then characterized independent T_2_ edited lines homozygous for different mutations and observed variable effects on plant height in both *Rht-B1a* (**Fig. 1a-c**) and *Rht-B1b* backgrounds (**Fig. S3a,b**), which correlated with the predicted effect of the mutations on miR172 binding. For example, in the *Rht-B1a* background the del6 mutation is predicted to be more detrimental to miR172 activity than del3 (**Fig. 1a**), and is associated with a larger reduction in plant height than del3 (**Fig. 1b,c**), even though none of them result in reading-frame shifts. The strongest *rAp2L-2* alleles in the *Rht-B1a* background reduced plant height to similar levels as *Rht-B1b* **(Fig. 1b,c**), and can be used to replace the *Rht1b* alleles. Taken together, these results indicate that CRISPR-Cas9 mutations in the miR172 target site of *Ap2l-B2* can be used to modulate wheat plant height, providing breeders flexibility to achieve their desired height targets.

The *rAp2l-B2* plants showed a higher spikelet density (**Fig. 1d, Fig. S3c)** as a result of reductions in spike length and slight increases in spikelet number per spike (**Data S4**). In the *Rth-B1a* background, the edited lines headed 1.8-2.9 days later, which was comparable to the delay generated by *Rht-B1b* (**Fig. 1e**). The delay in heading time associated with the *rAp2l-B2* alleles was slightly stronger in the *Rht-B1b* sister lines (4.4 to 5.7 days delay, **Fig. S3d, Data S4**).

Finally, plants with and without the *rAp2l-B2* mutations showed similar coleoptile and first-leaf lengths in both the *Rht-B1a* (**Fig. 1f,g, Data S4**) and *Rht-B1b* backgrounds (**Fig. S3e,f, Data S4**). In summary, these results indicate that the *rAp2l-2B* alleles can be used to reduce plant height with limited pleiotropic effects on spike architecture or heading time, and with beneficial effects in coleoptile length relative to the *Rht1b* alleles.

The highly efficient CRISPR vector makes it possible to rapidly induce different *rAp2l2* dwarfing alleles in elite backgrounds without time-consuming crosses. To demonstrate this strategy, we generated semi-dwarf mutants for the triticale cultivar ‘UC-Bopak’ (PVP 202100269). Triticale is an anthropogenic allohexaploid combining wheat and rye (AABBRR genomes), which delivers significantly high biomass and grain yield**^11^**. However, the taller plant stature of many triticale cultivars combined with their larger and heavier spikes can result in increased lodging. We transformed UC-Bopak using the same gRNA targeting the miR172 binding site in both *AP2L-B2* and *AP2L-R2* homeologs (**Fig. 1h**). Under greenhouse conditions, we observed a reduction in plant height in edited lines that correlated with the dosage of mutations and the predicted effect of the mutations on miR172 binding energy **(Fig. 1h,i, Fig. S4a,b)**. By selecting different combinations of *rAp2l2* mutations, we were able to fine-tune triticale plant height **(Fig. 1i, Fig. S4b)** without affecting coleoptile and first-leaf length or heading time **(Fig. 1j, Fig. S4c,e)**. The edited plants showed more compact spikes but with the same number of spikelets (**Fig. S4f, Data S5)**.

Finally, we evaluated lines with 1bp deletions in the miR172 target site of *AP2L-B2* (B) or *AP2L-R2* (R) under field conditions in two consecutive years. In 2023, we used headrows **(Fig. S5)** and in 2024 small yield plots as experimental units **(Fig. 1k-n)**. The plants with the 1-bp deletions were 17-18 cm shorter the first year (**Fig. S5a,b**) and 12-14 cm shorter the second year (**Fig. 1l**), indicating some interaction with the environment. The spikes of the edited lines were more compact than the wildtype (**Fig. S5c**), but this was not associated with significant differences in grain yield (**Fig. 1n, Fig. S5d**). The second year, the plots of the wildtype variety suffered significantly more lodging than the edited lines (*P*<0.0001, **Fig. 1k,m**). Although the differences in grain yield were not significant (**Fig. 1n, Fig. S5d**), the edited lines showed a combined 9.5% increase in grain yield the second year (*P=*0.0528, **Data S3**), which was likely associated with their superior resistance to lodging.

In summary, we demonstrate that different induced mutations in the miR172 target site of *AP2L2* genes can be used to precisely modulate wheat and triticale plant height. Breeders can use this technology to evaluate multiple plants heights in their top lines without lengthy backcrossing programs. Moreover, the *rAp2l2* alleles did not reduce coleoptile and first-leaf length, suggesting that they can be a valuable replacement of the gibberellin insensitive *Rht1b* alleles.

## Author contributions

**CZ** performed experiments and contributed to data analyses manuscript writing. **JH** contributed to project conceptualization, laboratory and field experiments, data analyses, and manuscript writing. **MP** performed experiments. **DT** conducted transformation experiments. **JD** contributed to project conceptualization, funding acquisition, CZ supervision, data analyses, and manuscript writing. **JMD** contributed to project conceptualization, performed and designed experiments, analyzed data, supervised MP, and wrote the original paper draft. **JMD** and **JD** generated the final version of the article.

## Acknowledgements

The authors acknowledge the support of UC Davis undergraduate students Chen Yang and Fangni Wu for their assistance in data collection, and UC Davis researcher Chengxia Li her revision of the manuscript.

## Data and germplasm availability

All the raw data and statistical analyses supporting all figures and supplemental figures are provided in the supplemental data Excel file (Data S1 to S6). Germplasm carrying the different mutations is available from the authors upon request and seeds are currently being increased to deposit them in GRIN-Global to facilitate long term public access.

## Online Methods

### Plant material and growing conditions

The tetraploid wheat variety Kronos used in this study has the *AP2L-5A* Q allele, which confers the subcompact spike phenotype and free-threshing character, and the *Rht-B1b* allele that confers the semi-dwarf phenotype. We also use a taller near-isogenic line of Kronos that was previously generated by introgressing the *Rht-B1a* allele from the tetraploid wheat cultivar ‘Gredho’ (PI 532239) into Kronos through six rounds of backcrossing (Kronos-*Rht-B1a*) ^1^. A KASP marker to differentiate the two *Rht-B1* alleles was also developed in the previous study ^1^.

Chemically induced mutants in the miR172 target site of the *AP2L2* genes were also described previously ^2^. These mutations reduce the ability of miR172 to cleave the *AP2L2* mRNAs and are designated as resistant alleles or *rAp2l2* alleles. The *rAp2l-A2* allele was previously generated in Kronos (mutant K2236) by ethyl methanesulfonate (EMS) ^2^, whereas the *rAp2l-B2* allele was generated with sodium azide in the winter hexaploid variety Wedgetail (*Rht-B1a* and *Rht-D1b*), which was mutagenized using sodium azide ^3^.

For this study, we combined the *rAp2l-A2* and *rAp2l-B2* mutations in Kronos by crossing the mutant lines Kronos-*rAp2l-A2* and Wedgetail-*rAp2l-B2*. As Wedgetail-*rAp2l-B2* is a winter cultivar, we vernalized it for seven weeks to induce flowering. The F_1_ plants were pollinated with Kronos-*rAp2l-A2*, and the resulting BC_1_F_1_ plants heterozygous for *rAp2l-A2* and *rAp2l-B2* were backcrossed to Kronos-*Rht-B1b* two more times. The BC_3_F_2_ population was genotyped using KAPS markers for *rAp2l-A2* and *rAp2l-B2* (primers in Table S1), and the four possible homozygous combinations were selected (wt, *rAp2l-A2, rAp2l-B2*, and *rAp2l-A2 rAp2l-B2*).

The commercially available triticale (× *Triticosecale*) cultivar used for transformation is a hexaploid (genomes AABBRR) spring cultivar from the UC Davis breeding program, released as ‘UC-Bopak’ (PVP 202100269). UC-Bopak is ∼110 cm tall and has very high grain yields relative to other triticales grown in the region.

Multiple independent transgenic plants were generated in Kronos, Kronos-*Rht-B1a* and UC-Bopak (see transformation and selection methods below). Transgenic plants in the T_0_ generation were genotyped by amplicon sequencing to identify edits in the miR172 target site of *AP2L2* genes. Selected T_0_ lines with edits in the target region were advanced to T_2_ in Kronos and to T_5_ in triticale. Plants were genotyped by amplicon sequencing to confirm the edits, and the presence of T-DNA sequences carrying the CRISPR-Cas9 was determined by PCR using primers described in Table S1. For triticale, we also crossed T_1_ plants with the wildtype UC-Bopak, and advanced the progeny to the F_3_ generation, where we selected transgene-free edited plants. Field experiments were performed in 2023 with F_4_ or T_5_ plants and in 2024 with F_5_ or T_6_ plants.

The transgenic plants were grown in cones in PGR15 growth chambers (Conviron, http://www.conviron.com) adjusted to 16 h of light (22°C) and 8 h of darkness (18°C). The intensity of the sodium halide lights measured at the height of plant heads was ∼260 μM m^-2^s^-1^.

For phenotypic evaluation, mutants and transgenic plants were grown in a greenhouse under long-day photoperiod (16-h light / 8-h dark, natural light supplemented with artificial light for 16 h). Temperatures oscillated between 23 °C during the day and 20 °C during the night. Plants were germinated in Petri dishes at 4°C for 3–5 days. After the first leaf emerged, we transplanted the seedlings into one-gallon pots (two plants each pot), and recorded days to heading from this day until emergency of half of the main spike from the flag leaf. Spikelet number per spike, spike length, and stem length (internodes and peduncle) measurements were taken at maturity.

### Field experiments

The CRISPR-edited lines of the triticale variety UC-Bopak with 1-bp deletions in the miR172 binding site of *AP2L-B2* (B del1) or *AP2L-R2* (R del1) and a sibling line without edits (wt) were evaluated in field experiments over two seasons at the UC Experimental Field Station in Davis, CA (38° 32′ N, 121° 46′ W). During both field seasons plants received a total of 225 kg/ha of N applied as ammonium sulfate, and irrigation and herbicide applications when needed. The 2023 field experiment was sown in January 2023 and harvested in June 2023. This experiment was organized in a completely randomized design using 2-m long rows as experimental units (∼100 grains per row) and consisted of 14 replications of wt, 15 replications of B del1 and 6 replications of R del1. Seeds harvested from these rows were used in field experiments in the 2024 field season that was sown in December 2023 and harvested in June 2024. The 2024 experiment included 8 replications for each genotype and was arranged in a completely randomized design, using small plots as experimental units (4.5 m x 1.4 m) with a seeding density of 262 grains per m^2^.

In both field seasons, we measured total plant height from the soil to the top of the main spike excluding awns. In the 2023 season, we randomly selected 4 to 6 primary tillers from each row, and obtained the length of peduncles, internodes, and spikes (in cm). We also determined the spikelet number per spike (SNS) and calculated spikelet density as SNS/spike length. Measurements from the 4-6 subsamples were averaged, and row means were used in the statistical analyses. We also determined grain yield per row (in grams) for the 2023 experiment. In the 2024 season, yield plots were evaluated for lodging on a 1-9 scale, where 1 indicates no lodging, 5 indicates more than half of the plot is lodged by 45 degrees or more, and 9 indicates the entire plot was flat on the ground. Grain from each yield plot was harvested with a Zurn 150 plot combine and yield was calculated based on measured plot length and converted to kg of grain per hectare.

### CRISPR vector and transformation

The miR172 target sites of the *AP2L-B2* in Kronos and both *AP2L-B2* and *AP2L-R2* in UC-Bopak were edited using CRISPR-Cas9. We designed a gRNA that specifically targets the miR172 sequence in these two homeologs (Fig. 1a, h). The gRNA was cloned into the binary vector JD633 that includes Cas9 and GRF4-GIF1 cassettes ^4^ (Addgene Plasmid #160393), and the final vector was transformed into the *Agrobacterium* strain EHA105. Transgenic plants were generated at the UC Davis Plant Transformation Facility (http://ucdptf.ucdavis.edu/). Immature embryos from Kronos-*Rht-B1a*, Kronos-*Rht-B1b* and triticale were inoculated with *Agrobacterium*. Transgenic plant selection was done using hygromycin, and transgene insertion was validated by DNA extraction and PCR.

### CRISPR Genotyping by amplicon sequencing

We collected leaf samples from plants and isolated genomic DNA using the commercial DNeasy kit (Qiagen), according to the manufacturer’s protocol. The detection of genome editing events was done by amplicon sequencing as described previously ^5^. Briefly, we performed polymerase chain reactions (PCRs) with primers flanking the regions targeted by the different gRNAs (primers in Table S1). Then, we added barcoded adaptors through a second nested PCR, pooled the PCR products, purified, and subjected them to CRISPR sequencing using the sequencing services provided by MGH CCIB DNA Core (https://dnacore.mgh.harvard.edu/new-cgi-bin/site/pages/crispr_sequencing_main.jsp)..

### Statistical analyses

The raw data and statistical analyses supporting the figures and supplemental figures in this study are provided in the Supplementary Data (excel file). Single way ANOVAS were performed for every trait. Homogeneity of variances were tested using Levene’s test and normality of residuals using the Shapiro-Wilk test. When ANOVA assumptions were not met, data was transformed using power transformations. In a few cases, when we could not find any appropriate transformation to satisfy both ANOVA assumptions, we used pairwise non-parametric Kruskal-Wallis tests. All statistical analyses were performed using SAS version 9.4.

Distribution of the data is presented using box-plots except for stem length where bar graphs were used to present the individual contributions of the peduncle and internodes to the total stem length. In the box-plots (generated with Excel), the middle line of the box represents the median and the x represents the mean. The bottom line of the box represents the first quartile and the top line the third quartile. The whiskers extend from the ends of the box to the minimum and maximum values.

To calculate the degree of dominance of the *rAp2l2* alleles, we genotyped segregating T_0_ and T_1_ lines in the *Rht-B1a* and *Rht-B1b* backgrounds and classified the plants as heterozygous, homozygous wildtype or homozygous/biallelic mutant. Degree of dominance was calculated as the difference between the average stem length of the heterozygotes and the midpoint value between the homozygotes, divided by the additive effect ^6^.

## Supplementary Figures

**Figure S1.**
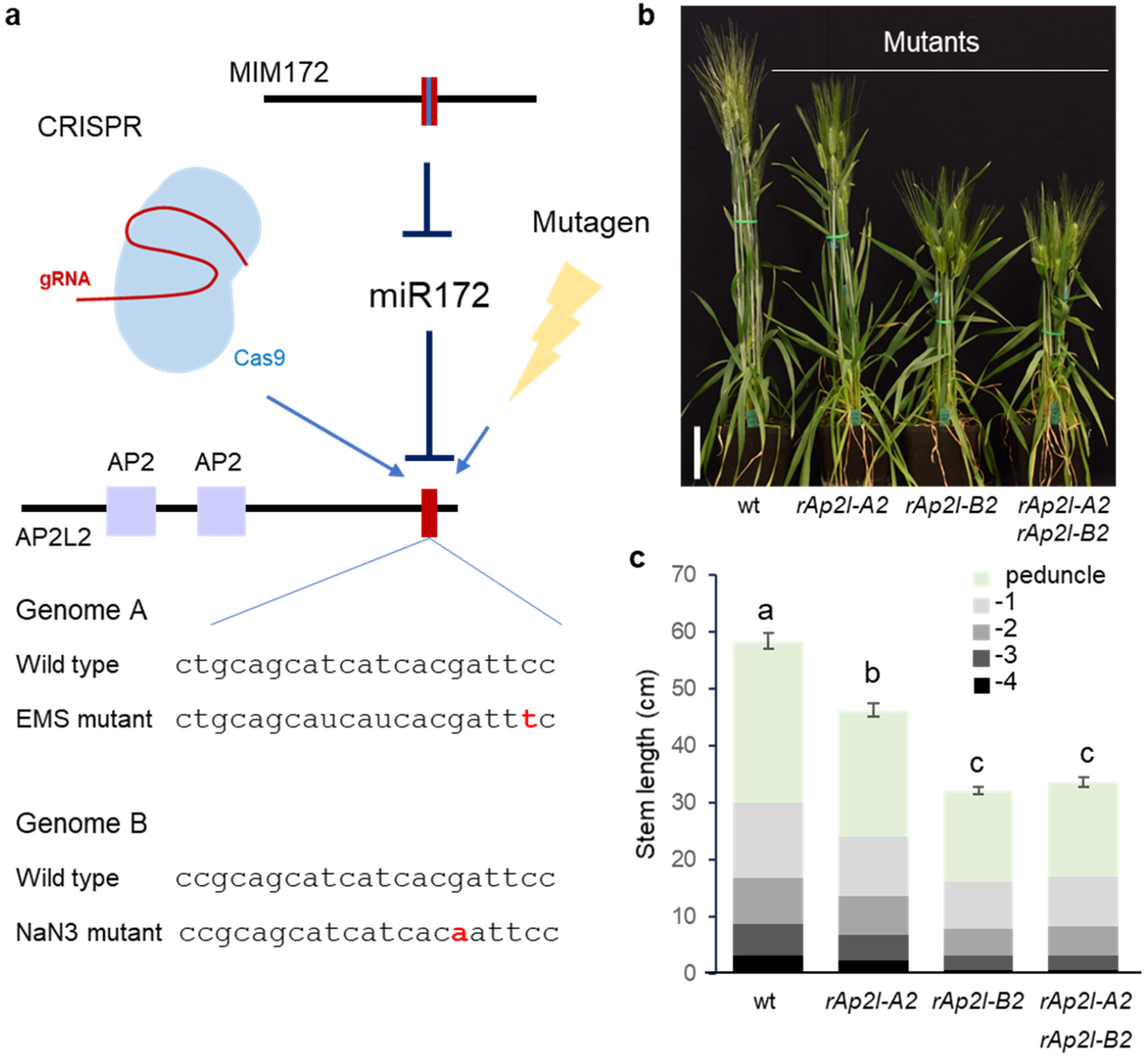
Reduced miR172 activity on *AP2L2* results in shorter plants. **a**. Different approaches to modulate miR172 - *AP2L2* interaction. Mature miR172 activity can be reduced by expressing a target mimicry (MIM172), or by mutating the miR172 target site in *AP2L* genes by CRISPR or chemical mutagenesis. The regions encoding the AP2 domains are indicated in purple and the miR172 target site is in red. Below are the *AP2L-A2* and *AP2L-B2* miR172 target sites in wildtype and *rAp2l2* alleles generated by chemical mutagenesis (in red). **b**. Selected plants three weeks after heading carrying wildtype, *rAp2l-A2, rAp2l-B2*, and combined mutant alleles. Bar= 10 cm. **c**. Stem length: internodes are in different gray colors and peduncles are in green (−1 is the closest internode to the peduncle). Different letters above the plots indicate significant differences in total stem length based on Tukey tests (*P*<0.05, n= 16 plants per genotype). These experiments are in a Kronos semi-dwarf background (*Rht-B1b*). Raw data and statistics are available in Data S1.

**Figure S2.**
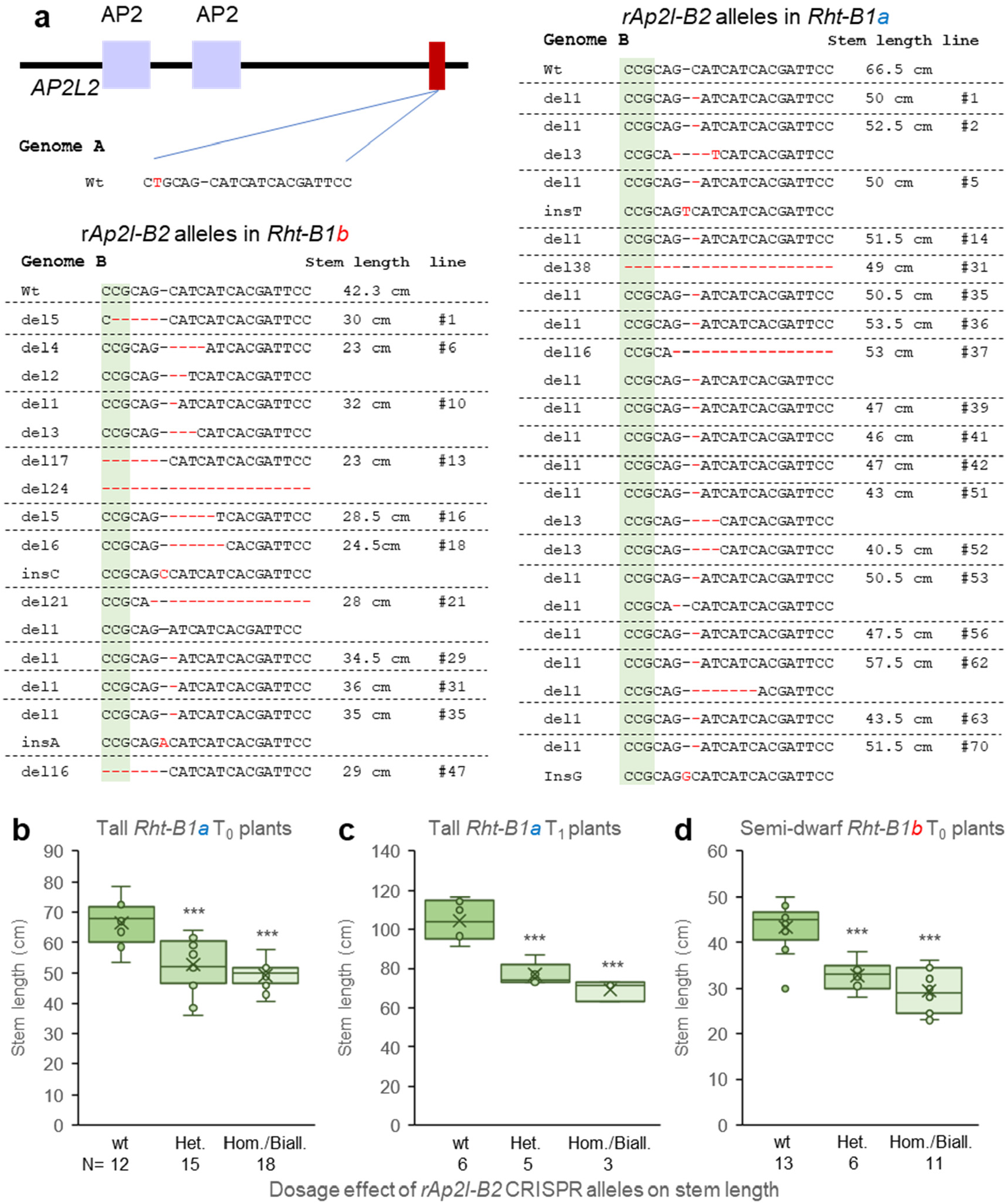
CRISPR induced *rAp2l-B2* alleles show a semidominant dwarfing effect. **a**. Schematic representation of the miR172 target sites and different *rAp2l-B2* CRISPR alleles from independent T_0_ lines in semi-dwarf Kronos-*Rht-B1b* and its tall sister line Kronos-*Rht-B1a* (del= deletion, ins= insertion). Stem length is indicated in the right column. (**b-d**) Dosage effect of *rAp2l-B2* CRISPR alleles on stem length (wt= wildtype, Het= heterozygous, Hom./Biall= homozygous or biallelic based). (**b**) Forty-five independent T_0_ events in tall Kronos-*Rht-B1a* grown in small cones. (**c)** Fourteen T_1_ progeny of a T_0_ line heterozygous for del6 in the *Rht-B1a* background grown in larger pots. (**d)** Thirty independent T_0_ lines in the Kronos-*Rht-B1b* background grown in small cones. *** = *P*<0.001 based on Dunnett tests against the wildtype. Numbers below genotypes indicate number of plants tested for each genotype. Raw data and statistics are available in Data S2.

**Figure S3.**
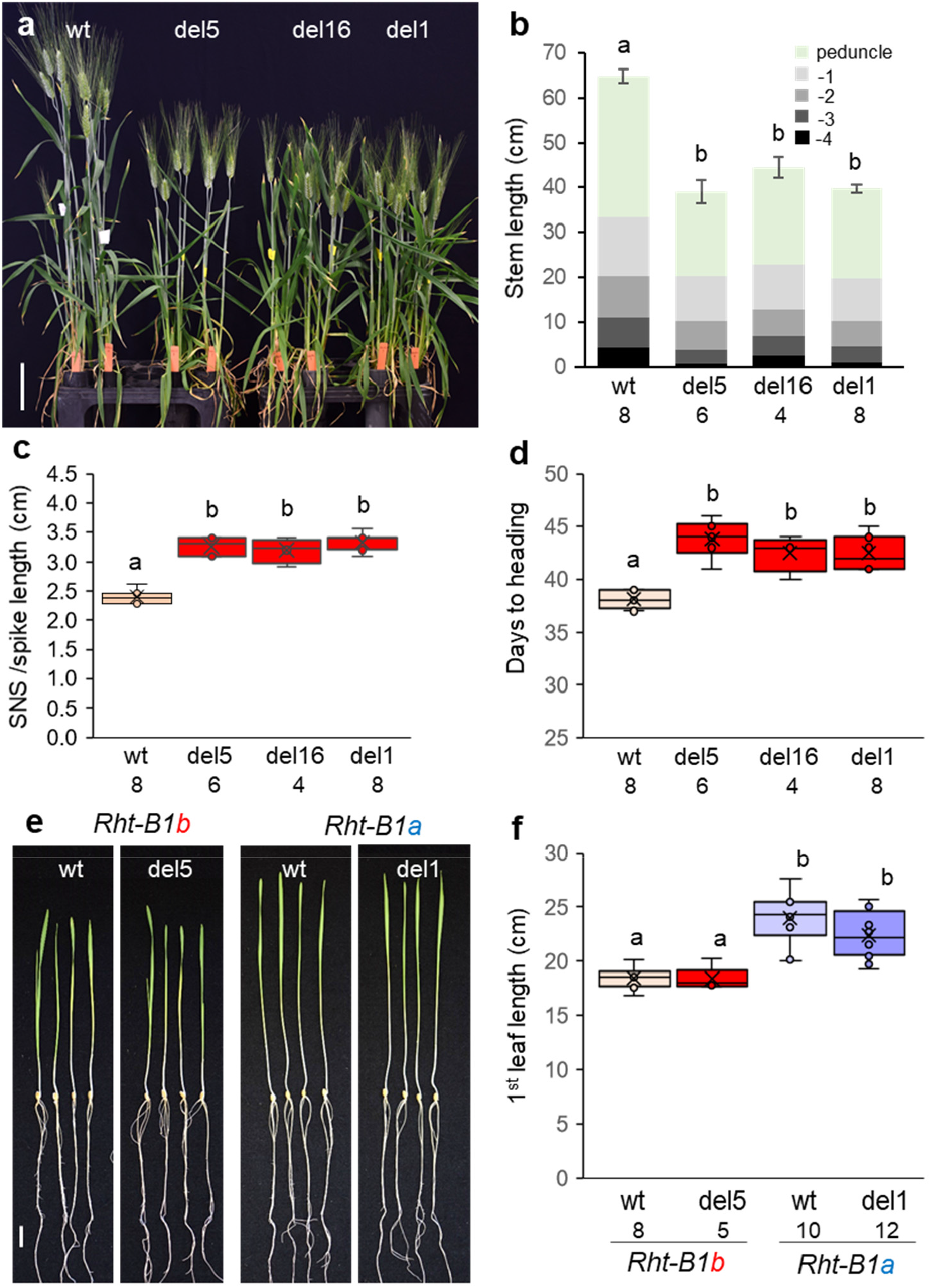
Effect of CRISPR-edits in the miR172 binding site of *AP2L-B2*. (**a**) Selected Kronos plants after heading carrying the wildtype allele and three independent CRISPR-Cas9 induced deletions. Bar = 10 cm. (**b**) Stem length: internodes are in different gray colors and peduncles are in green (n= 16 plants per genotype). (**c**) Spikelet density calculated as the ratio between spikelet number per spike (SNS) and spike length in cm. (**d**) Days to heading. **a** to **d** in Kronos-*Rht-B1b* background. (**e**) Seedlings of wildtype (wt) and edited lines in *Rht-B1b* and *Rht-B1a* backgrounds. Bar = 2 cm. (**f**) Length of the first leaf. Note the shorter seedlings in *Rht-B1b* relative to *Rht-B1a*. Different letters above the bars or box-plots indicate significant differences based on Tukey tests (*P*<0.05). Numbers below the genotypes indicate the number of plants measured per genotype. Raw data and statistics (including coleoptile length) are available in Data S4.

**Figure S4.**
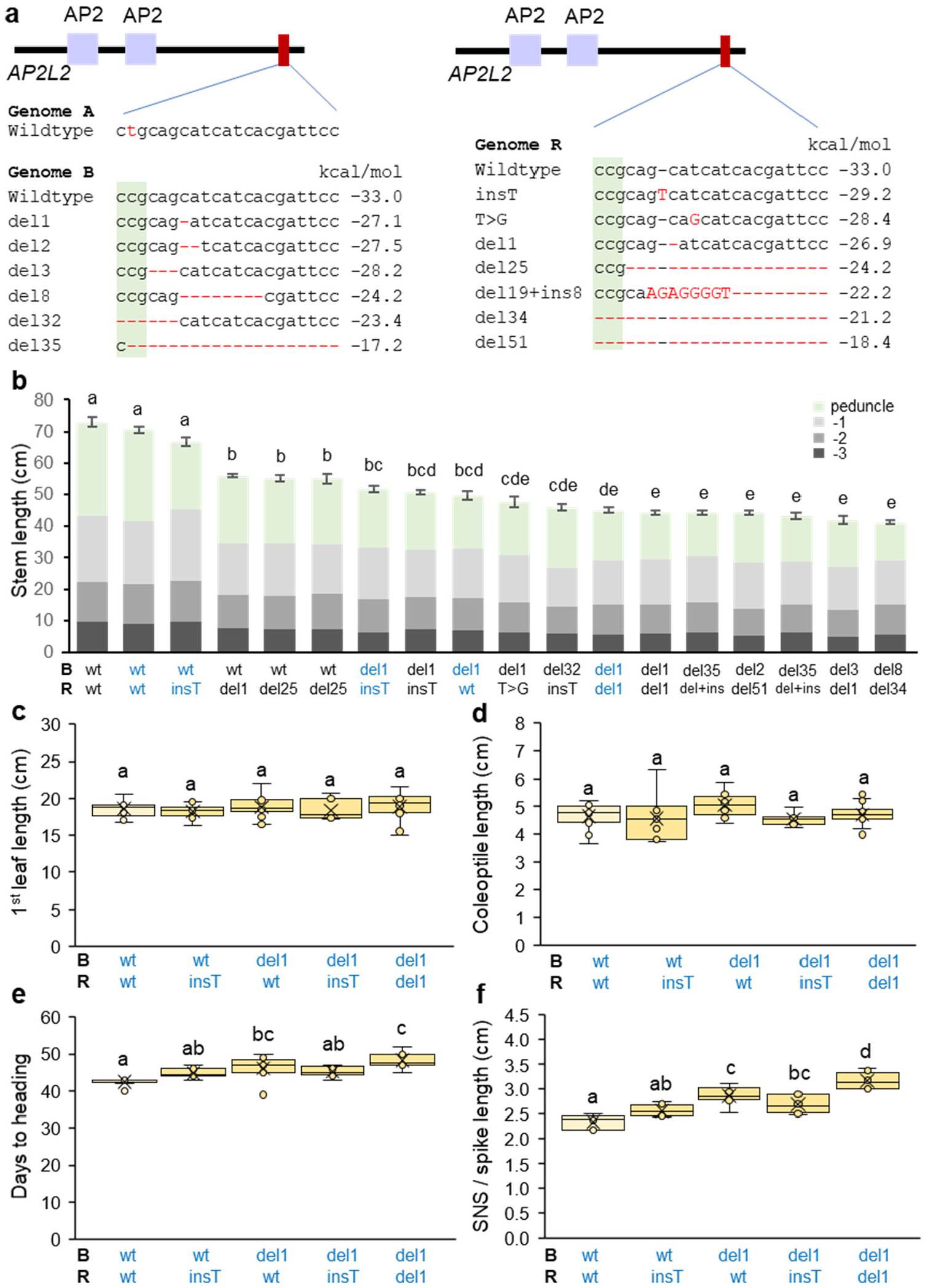
*rAp2l-B2* and *rAp2l-R2* alleles modulate plant height in triticale cultivar UC-Bopak. a. Schematic representation of *AP2L2* and sequences of the CRISPR-edited sites in the miR172 target sites in *AP2L-B2* (B genome from wheat) and *AP2L-R2* (R genome from rye). del= deletion, ins= insertion. The estimated interaction energy in kcal/mol is indicated to the right. **b**. Stem length in selected lines: internodes are in different grey colors and peduncles are in green (n=8 except wt/del25 n=6). The *AP2L2* genotypes of the B and R genomes are indicate below each bar. Genotypes in blue are the same in panels **b** to **f**. (**c**) Length of first leaf in 10 days-old seedlings. (**d**) Coleoptile length in cm. (**e**) Days to heading. (**f**) Spikelet density (spikelet number per spike / spike length in cm). **c-f** n=8. Different letters above bars and plots indicate significant differences based on Tukey tests (*P*<0.05). Raw data and statistics are available in Data S5.

**Figure S5.**
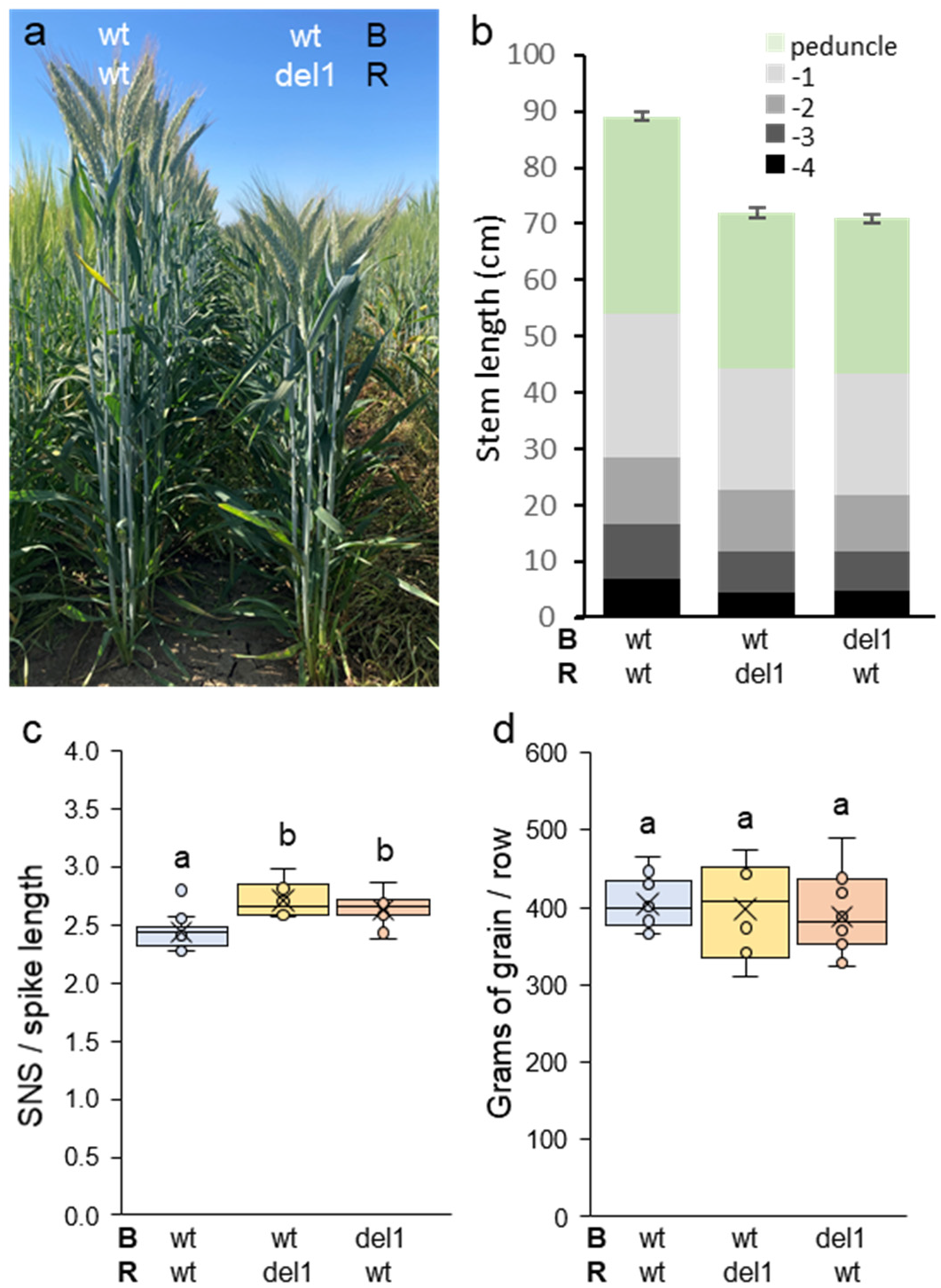
Field experiment (2023 field season) comparing the triticale variety UC-Bopak and CRISPR lines with 1-bp deletions in the miR172 binding site of *AP2L-B2* (wheat B genome) or *AP2L-R2* (rye R genome). **a**. Selected rows comparing UC-Bopak with a CRISPR-edited line carrying the 1-bp deletion in *AP2L-R2*. **b**. Stem length: internodes are in different gray colors and peduncles are in green. **c**. Spikelet density (spikelet number per spike / spike length in cm). **d**. Grams of grains produced per row. wt/wt= 14 headrows, wt/del1= 6 headrows, del1/wt= 15 headrows. Headrows were used as experimental unites in a completely randomized design. Four to six plants (subsamples) were measured per row and values were averaged for the statistical analyses. Different letters above the bars and box-plots indicate significant differences based on Tukey tests (*P*<0.05). Raw data and statistics are available in Data S6.

## Supplementary Table

**Table S1.**
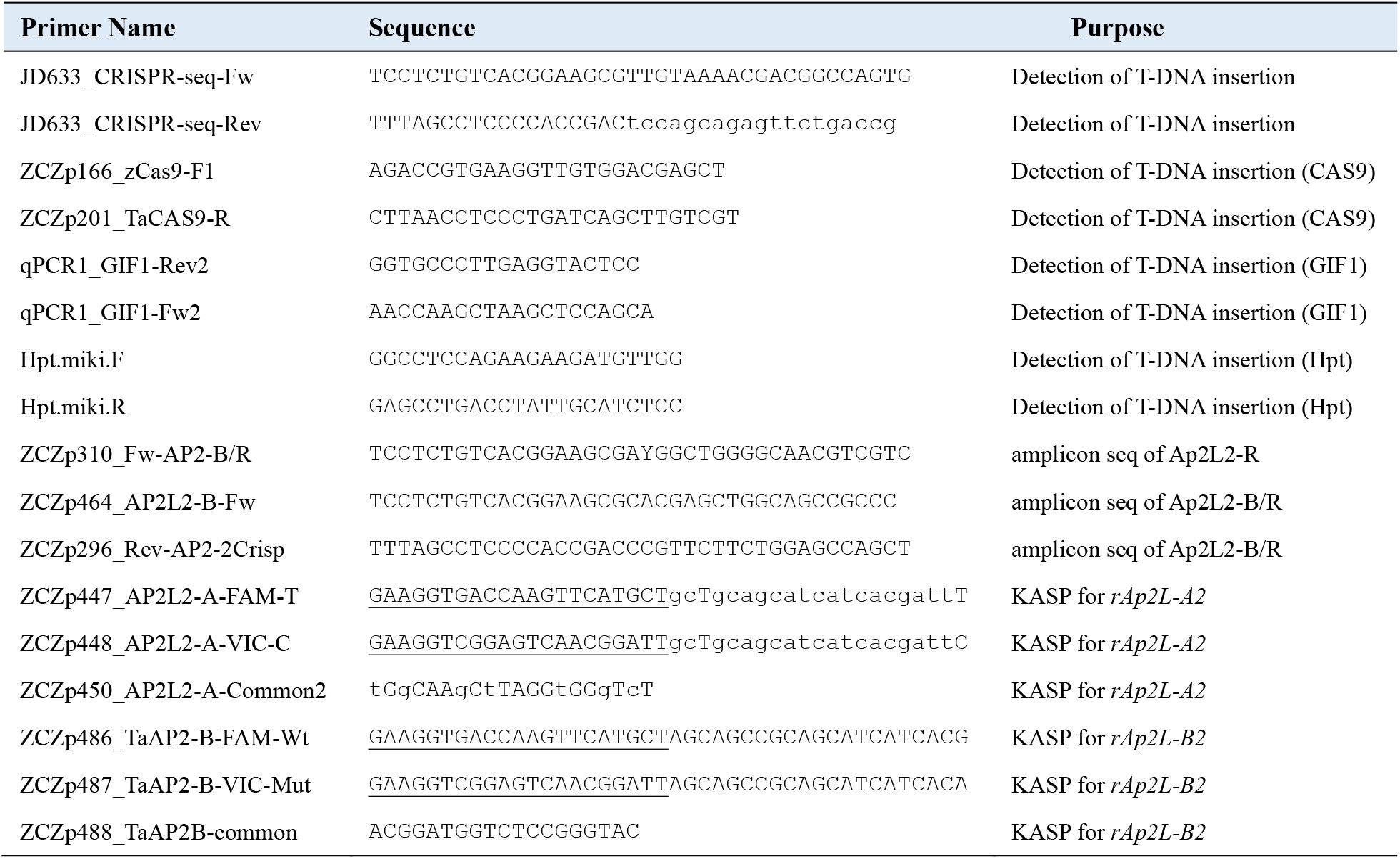
Primers used in this study.

